# mgPGPT: Metagenomic analysis of plant growth-promoting traits

**DOI:** 10.1101/2024.02.17.580828

**Authors:** Sascha Patz, Mario Rauh, Anupam Gautam, Daniel H. Huson

**Author notes:** Address correspondence to Sascha Patz,.

## Abstract

In a recent publication, we introduce the PGPT ontology and PGPT-db of bacterial plant growth-promotion traits and associated database of protein sequences, and provide several tools for bacterial genome analysis on the PLaBAse server. Here, we extend the scope of the PGPT ontology to perform PGPT analysis of metagenomic datasets. First, we introduce mgPGPT-db, an extended database of 39, 582, 183 protein sequences obtained computationally by including proteins from AnnoTree. With this, we have integrated the PGPT ontology into our metagenome analysis tool MEGAN and provide mapping files to identify PGPT-related genes using the results of a DIAMOND alignment of reads against either the new mgPGPT-db database, the NCBI-nr protein database, or the AnnoTree protein database. We demonstrate and compare these different approaches in detail on an example data set and evince the improvement compared to the PGPT-db. We also compare the inferred PGPT content of several samples taken from different environments and reveal plant specific PGPT clustering.

**IMPORTANCE:** A deeper understanding of plant growth-promoting traits of bacteria is important to enlight and enhance the native plant-beneficial bacterial functional diversity regarding the environmental stress adaption or the ability to suppress even food-borne pathogens by strain inoculation dedicated to agriculture and other plant production systems. The work presented here extends recent work beyond the analysis of individual to allow the assessment of the PGPT potential of metagenomes obtained from environmental samples.

## INTRODUCTION

Microbes with plant growth promoting traits (PGPTs) are considered potential bioinoculants whose application to plant production systems may achieve beneficial effects such as plant growth-enhancement, plant stress-reduction or phytosanitary effects (1, 2). At present, only a small number of such plant growth-promoting bacteria and fungi (PGPBs/PGPFs) are used in agriculture (3, 4).

The in-silico detection and characterization of beneficial strains has gained a lot of importance (5, 6, 7). Recent approaches target the of synthetic synergistic communities (SynComs), obtained e.g., from one or multiple plant microbiomes that show best growth conditions even under biotic or abiotic stress, aiming at a broader target range, higher robustness and a better plant-beneficial efficacy (8, 9). The interaction between the microbiome and its host is poorly understood. In particular, the impact of the host genotype, native microbiome, metabolome, and features for differentiation between “friend and foe”, as well as the effect of environmental parameters on the plant growth, are unclear (10, 11, 12, 13).

One of the reason could be found in the fact, that until recently the annotation of plant growth-promoting traits (PGPTs), involved in such beneficial processes of putative bioinoculant strains, was still challenging and mostly manually curated resulting in non-standardized and less-comparable data sets (14, 15). Respective functional annotations were often obtained from EggNOG/COG (16) or KEGG (17) annotations and individually picked (18, 19, 20, 21). Likewise, metagenomic approaches just addressed a arbitrary selection of important PGPTs by searching respective functional classes or identifiers (22, 23). These were often related to e.g., phytohormone production, nutrient acquisition, bioremediation or stress tolerance (24, 25).

Here, for instance, the metagenome analysis pipeline DIAMOND+MEGAN6 (26, 27, 28, 29, 30) can be used, as it comes up with the conservative read alignment approach against NCBInr proteins that are further mapped to NCBI (31) or GTDB (32) taxonomy and various functional classifications, including KEGG. For example, Nisrina and coworkers applied a MEGAN approach for PGPT screening of PGPBs in fungi suppressive soils (33). Recently, the AnnoTree assignment approach was added to MEGAN, that showed higher accuracy especially for environmental samples like soil, addressing a huge amount of uncultured strains as it relies on additional protein sequences derived from metagenome-assembled-genomes (MAGS) (34, 35). Consequently, the MEGAN NCBInr database was extended to include these additional sequences (release February 2022). But, as said, both approaches only allow the manual screening for plant-beneficial traits. The only established standardized PGPT annotation approach is currently available via PLaBAse as tool PGPT-Pred for genomic protein-to-PGPT annotations of single bacterial strains. (36).

In the present study, we have (i) implemented the PGPT ontology into MEGAN6 as PGPT viewer, (ii) established a mgPGPT protein and mapping database, and (iii) extended the existent NCBInr and Annotree mapping databases by PGPT annoatations. Hence, we provide three metagenomic PGPT analysis pipelines for short and long read assignments to PGPTs. The mgPGPT protein database extends the current available PGPT-db on PLaBAse, as it comprises even metagenomic protein sequences from e.g., uncultivated strains. Finally, our performance tests on a single data set and a MWAS study evince the improved annotation approach for mgPGPTs and highlight the impact of the PGPT ontology for understanding the PGPT variability among metagenomes associated not only with the environment plant (phyllosphere and rhizosphere) but also to soil, human, and animal to be studied in detail in the future.

## RESULTS

The DIAMOND+MEGAN pipeline is appropriate for taxonomic and functional classification of metagenomic samples. Here, we aim to extend its functional capability by annotation of PGPTs for environmental samples to mainly classify plant- and soil-associated bacterial metagenomes. We provide three major approaches to perform PGPT analysis, each starting with the diamond read alignment against a respective protein database followed by meganization against a related mapping database, and classification and visualization with MEGAN (Figure 1).

**FIG 1.**
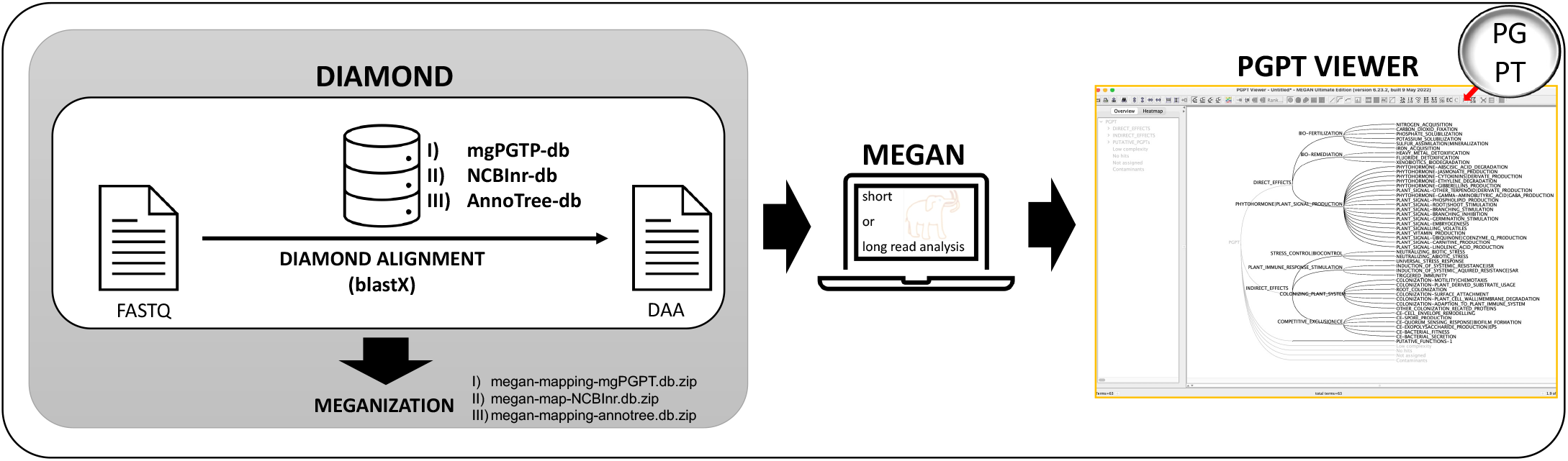
Metagenomic Analysis of PGPTs using MEGAN6. DIAMOND+MEGAN pipline for PGPT mapping via three different approaches: (i) the conventional NCBInr -or (ii) the AnnoTree-db based read mappings, and (iii) the mgPGPT-db mapping.

### mgPGPT-db based metagenomic PGPT analysis

The mgPGPT-db constitutes a protein database of PGPT-related proteins that is enriched in sequences achieved the AnnoTree-db database that includes proteins from metagenomic assembled genomes (MAGs) as well. The repsecitve mapping database stores to each PGPT protein identifier a taxonomic node id and a PGPT ontology node id needed for MEGAN analysis. Thus, the here established approach addresses the metagenomic functional analyses of PGPTs by read alignment and assignment against the mgPGPT-db and mgPGPT mapping database, available under https://plabase.cs.uni-tuebingen.de/pb/download.php. It allows direct PGPT assignment of reads and their taxonomic affiliation, but ignores taxonomic and functional extent of non-PGPT assigend reads.

### NCBInr-db based metagenomic PGPT analysis

The second approach dedicates the holisitc taxonomic and functional classification of all aligned reads including the PGPT content. It makes use of the conventional NCBInr protein and mapping databases for MEGAN analysis. To do so, the recent NCBInr mapping database for download includes a column for protein PGPT assignments that enables the autamitic PGPT content analysis. As that modification adresses only the read mapping step, older NCBInr based read alignemnt files can easily be meganized again using the new mapping database provided at https://software-ab.cs.uni-tuebingen.de/download/megan6/welcome.html

### AnnoTree-db based metagenomic PGPT analysis

Recently, our group provides the opportunity to use AnnoTree-based annotations for holisitic metagenomic analysis with MEGAN. Accordingly, we provide an extension of the AnnoTree mapping database for MEGAN, that includes PGPT assignments. It is avaialable at https://software-ab.cs.uni-tuebingen.de/download/megan-annotree/welcome.html

### PGPT ontology implementation in MEGAN6

For classification and visualization of PGPT-assigned reads the PGPT ontology with its 6,912 PGPTs is implemented in MEGAN6 as PGPT viewer. Thus, it allows analysis of short or long read alignments. As for some PGPTs up to 13 different functionalities appear the tree is multilabeled according to multiple functional roles a PGPT might achieve.

### Comparison of PGPT-assigned protein database contents

The mgPGPT-db comprises 39,582,183 protein sequences (Figure 2 A), means 5 times more sequences than the PGPT-db database, that was actually established for genomic protein annotation with the PGPT-Pred tool. Among all sequences 36,767,435 mgPGPT protein sequences were found in the AnnoTree (total protein count of 106,052,079) and 30,313,055 in NCBInr (346,290,401) protein databases. As orientation, KEGG annotated proteins of the mentioned databases range from around 52 (NCBInr-db) to around 57 million (AnnoTree-db). In total the overlap of the mgPGPT sequences to both databases sums up to 38,996,170 protein sequences, indicating a higher proportion of representative PGPT proteins covered by the mgPGPT-db including a remainder of additionally 586,013 proteins (Figure 2 B).

**FIG 2.**
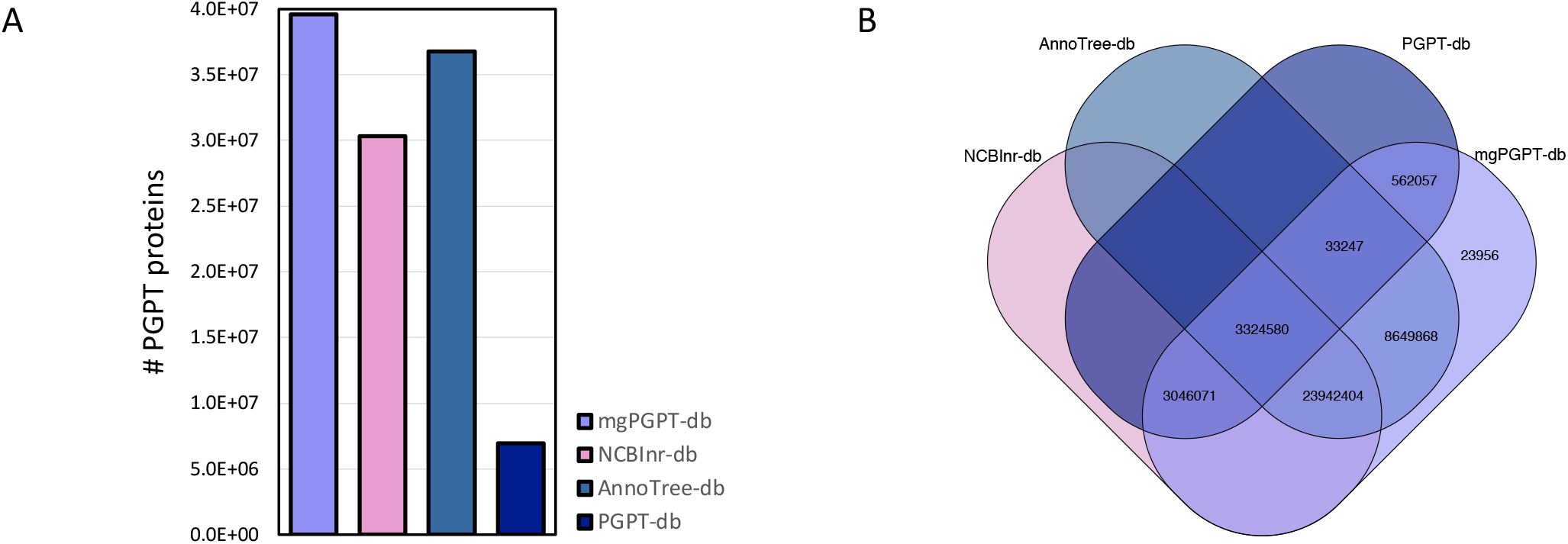
Overview about the PGPT content of all protein databases applicable via the DIAMOND+MEGAN Pipeline. (A) PGPT protein count among the metagenomic protein databases (i) mgPGPT-db, (ii) NCBInr-db, and (iii) AnnoTree-db compared to the original PGPT-db that is applied to genomic analysis. (B) Overlaps of PGPT-assigned proteins between respective databases.

### Performance comparison of PGPT assessment by all three metagenomic PGPT approaches

As an example, we used the metagenome sample mgm4626743.3, the mgPGPT-db based approach achieved highest read-to-PGPT assignment rates for all level 0 and level 1 nodes in the PGPT ontology, followed closely by the AnnoTree-db based PGPT mapping (Table 1). The NCBInr-db approach still shows adequate assignments rates. Accordingly, the PGPT-based metagenomic clustering (Figure 3 A,C), showed highest similarity between the mgPGPT- and AnnoTree-db approach with very low variance alterations on PC2. The original PGPT-db, usually dedicated to genomic protein sequences and here applied for comparison, lag behind all strategies in terms of PGPT assignments and PGPT functional clustering. When comparing all three metagenomic approaches, PGPT annotation based on the mgPGPT-db results in 8.624 million PGPT assigened reads, using AnnoTree-db in 8.279 million reads, and using NCBInr-db in 7.147 million reads (Table S1). While mgPGPT-db vs. AnnoTree-db assignments show an overlapp of 8,278,185 reads with a similarity of PGPT id assignments of 98% among them, the mgPGPT-db vs. NCBInr-db approach shares only 7,146,644 PGPT assigned reads with 95% similarity in the PGPT id assignments. All approaches together approximate 94% similarity. Among all three metagenomic approaches, the runtime was lowest for the mgPGPT approach followed by AnnoTree-db and then by NCBInr-db (Table S2).

**TABLE 1.** Performance of multiple PGPT annotation approaches on a single metagenomic dataset. Absolute read counts for PGPT ontology level 0 and 1 achieved by NCBI-db, AnnoTree-db, PGPT-db- or mgPGPT-db read mapping of sample mgm4626743.3.^*a*^.

| Protein Database: | mgPGPT-db<sup>b</sup> | NCBI-db<sup>b</sup> | AnnoTree-db<sup>b</sup> | PGPT-db<sup>c</sup> |
| --- | --- | --- | --- | --- |
| Total read count |  | 16,538,321 |  |  |
| Total aligned reads | 9,210,098 | **16,537,915** | *16,155,154* | 6,648,014 |
| Total assigned reads to NCBI taxonomy | 8,354,952 | **16,382,865** | *15,548,829* | 4,939,683 |
| Total assigned reads to PGPT-Level 0 | **8,624,177** | 7,147,454 | *8,279,000* | 5,109,800 |
| ... to PGPT-Level 1: Direct effects | **3,584,751** | 2,973,053 | *3,432,563* | 2,093,238 |
| ... to PGPT-Level 1: Indirect effects | **7,185,114** | 5,981,471 | *6,911,528* | 4,302,333 |
| ... to PGPT-Level 1: Putative PGPTs | **48,745** | 34,993 | *43,167* | 28,806 |

<sup>a</sup>Maximal counts per row are marked in bold and 2nd highest counts in italic. Interesting values are highlighted yellow.
<sup>b</sup>Available for DIAMOND+MEGAN pipeline
<sup>c</sup>Available via PGPT-Pred tool on PLABase

**FIG 3.**
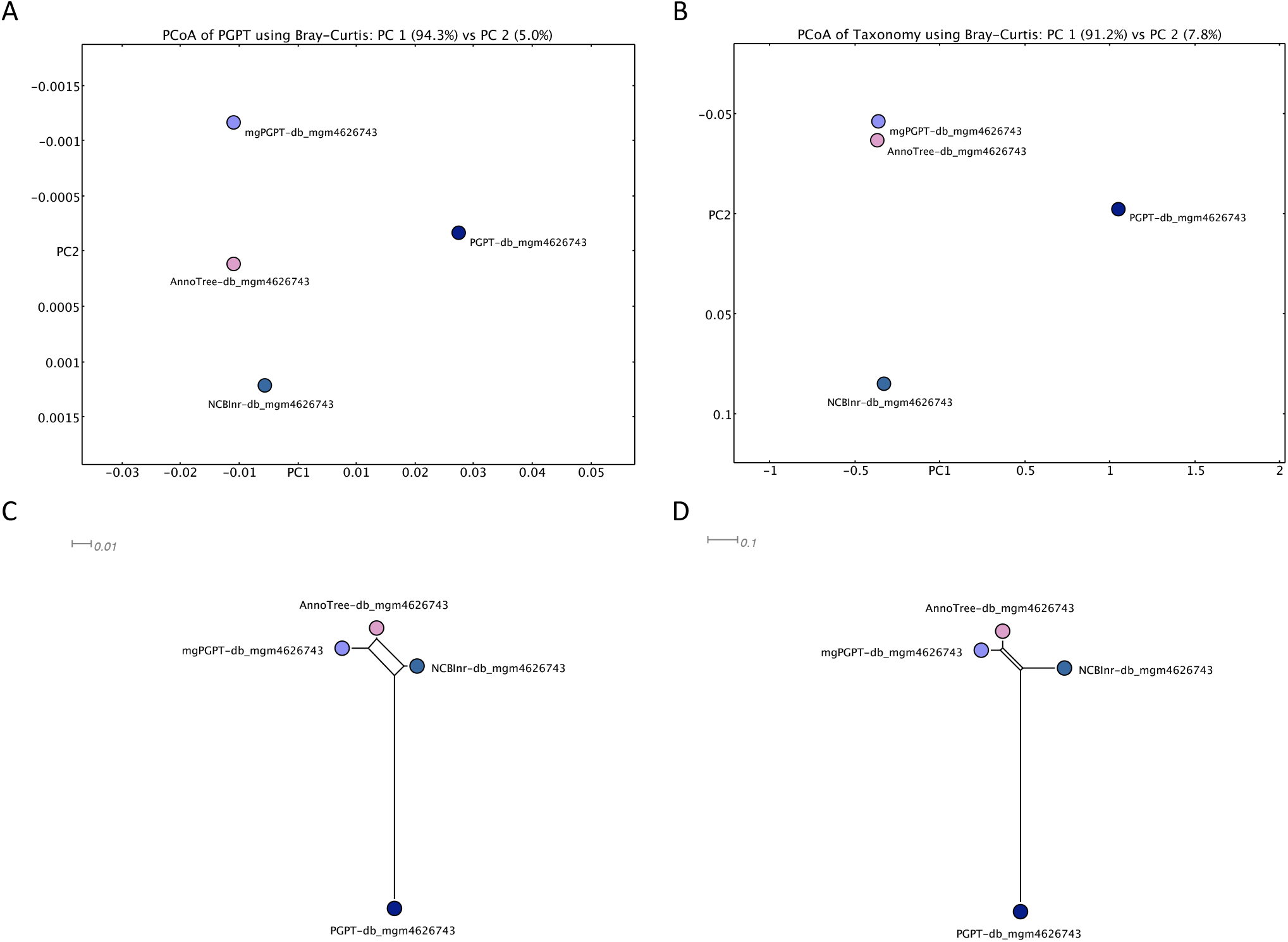
Single metagenome analysis comparison performed with all three metagenomic PGPT annotation approaches and the PGPT-db based PGPT mapping as reference. (A,C) Bray-Curtis PCoA and Outline plot for PGPT functional analysis on PGPT ontology level 6 reveals high comparability for all metagenomic approaches; (B,D) PCoA and Outline plot of the taxonomic analysis on Species level indicates best comparability between mgPGPT-db and AnnoTree-db based analyses. Metagenomic comparison is based on MEGAN6 normalization to lowest aligned read count found among all samples.

### Performance comparison of the taxonomic assessment by all three metagenomic PGPT approaches

Regarding taxonomic assignments we conclude, that with decreasing database size the taxonomic assigned read count declines along the decreasing rate of aligned reads (Table 1). Nethertheless, we evince highest similarity between the mgPGPT- and AnnoTree-db-based taxonomic clustering on species level, even by highly different taxonomic assigned read counts (Figure 3 B,D). When comparing the taxonomic affiliation of PGPT assigned reads only, we get, under the applied alignment and meganization parameters, for mgPGPT vs. NCBI around 51% and for mgPGPT vs. AnnoTree around 67% identical LCA assignments (Table S1). The rate of identical taxonomic hits for AnnoTree vs. NCBI is of around 62%.

### Improvement of the metagenomic PGPT appraoches compared to the previous PGPT-db approach

When applying the recent PGPT-db, with approx. 9.6 million PGPT assigend proteins, to metagenomics we achieve approximately 1 to 3.5 million read assignments less than using any of the three metagenomic approaches. The mgPGPT-db and AnnoTree-db based methods showed the highest difference in the PGPT assigment rate on level 0 and level 1.

### Performance comparison on a MWAS study

When analyzing and comparing multiple metagenomic samples across various environments, we get quite similar patterns for read assignment and alignment rates as for a single sample, considering all metagenomic PGPT approaches. Again, the entire read assignment rate for all nodes of the PGPT ontology level 0 and level 1 is best of mgPGPT-db followed by AnnoTree-db and then by NCBInr-db PGPT mappings (Table 2).

**TABLE 2.**
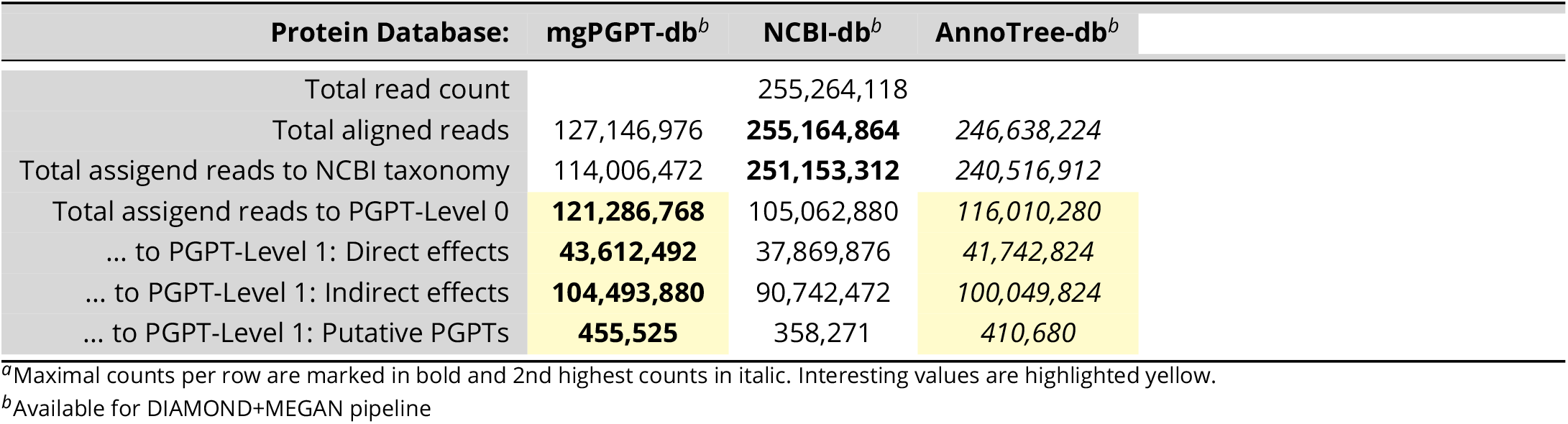
Summarized read assignment statistics of a MWAS study comprising 67 metagenomic samples of various environments. Shown are additive absolute read counts achieved by alignment against the respective PGPT-assigned protein databases.^*a*^.

| Protein Database: | mgPGPT-db<sup>b</sup> | NCBI-db<sup>b</sup> | AnnoTree-db<sup>b</sup> |
| --- | --- | --- | --- |
| Total read count |  | 255,264,118 |  |
| Total aligned reads | 127,146,976 | **255,164,864** | 246,638,224 |
| Total assigned reads to NCBI taxonomy | 114,006,472 | **251,153,312** | 240,516,912 |
| Total assigned reads to PGPT-Level 0 | **121,286,768** | 105,062,880 | *116,010,280* |
| ... to PGPT-Level 1: Direct effects | **43,612,492** | 37,869,876 | *41,742,824* |
| ... to PGPT-Level 1: Indirect effects | **104,493,880** | 90,742,472 | *100,049,824* |
| ... to PGPT-Level 1: Putative PGPTs | **455,525** | 358,271 | *410,680* |

<sup>a</sup>Maximal counts per row are marked in bold and 2nd highest counts in italic. Interesting values are highlighted yellow.<sup>b</sup>Available for DIAMOND+MEGAN pipeline

In accordance with the description of the single sample, MEGAN metagenomic functional clustering based on PGPTs at leaf level 6 shows similar results for all approaches (Figure 4 A-C). Even if using the smaller database mgPGPT-db we achieve comparable values of explained variance between samples from different environments. All approaches point to the fact, that plant-associated microbiomes analyzed here have distinct PGPT patterns with highest interference between rhizospheric and soil samples in contrast to the distinct clusters for human and animal samples. Two rhizospheric samples, mgm4778843.3 and mgm4732622.3 seem to cluster together with phyllospheric samples, but build an own cluster in the PC3 dimension. Both samples were isolated from Iranian soil enriched with iron ore. Likewise, one human sample seems to cluster closer to phyllospheric samples also show distinct clustering towards dimension PC3. It originate from the gut site and represent a sole site among the human samples origins.

**FIG 4.**
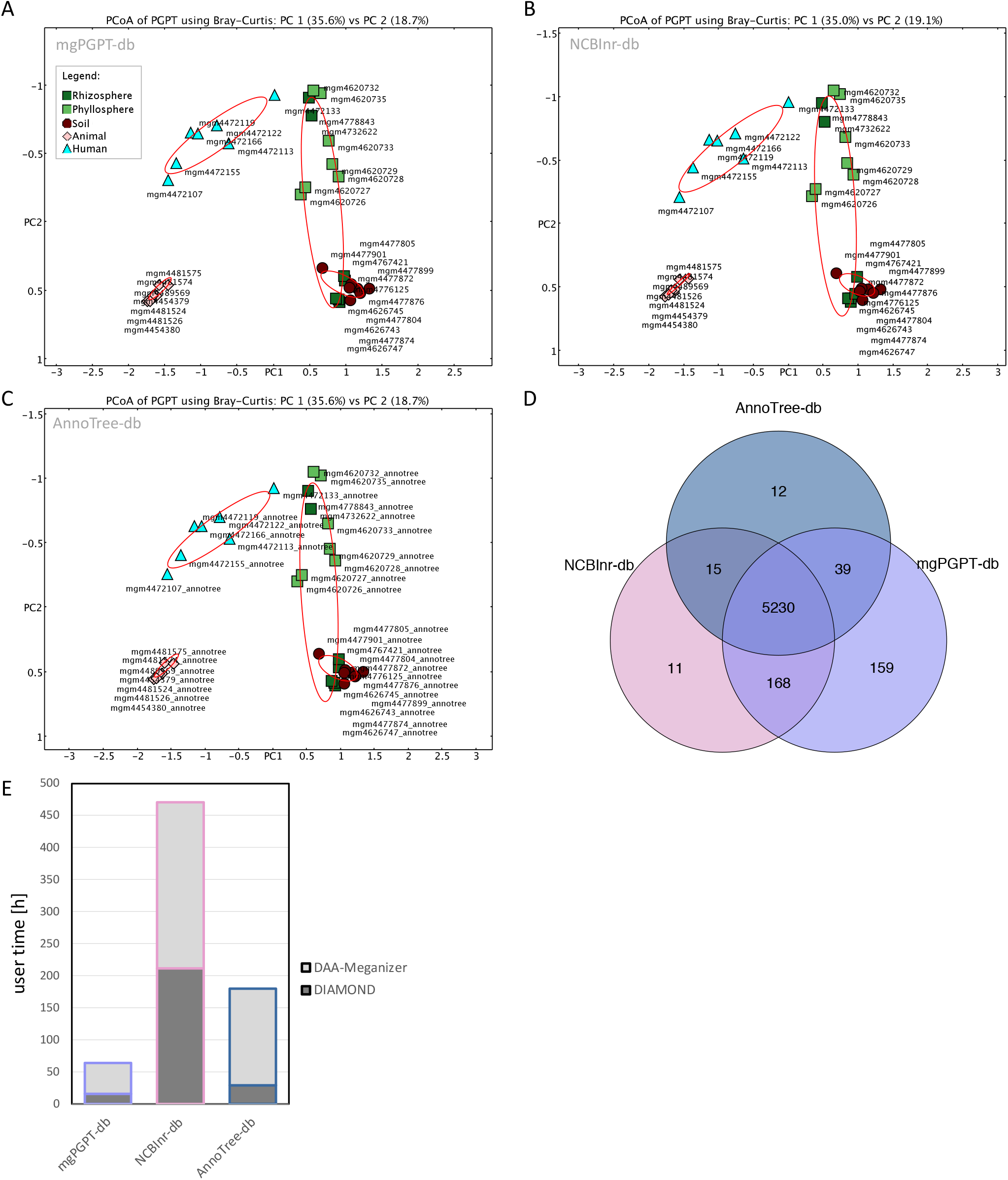
Metagenome-wide association study comprising five environmental classes performed with all three metagenomic PGPT annotation approaches using either mgPGPT-, NCBInr- or AnnoTree-db. (A-C) Bray-Curtis PCoA clustering of PGPT abundancies on PGPT ontology level 6 for the environmental classes plant-phyllosphere (green), plant-rhizosphere (darkgreen), human- (lightblue), animal- (coral) and soil-associated (brown) achieved by the above indicated metagenomic approaches. All values were normalized to the lowest aligned read count found among the samples. Red circles describe the respective groups: plant-, soil-, human- and animal-associated. (D) Overlap of detected PGPT ids by all approaches summarized as Venn diagram. (E) Runtime analysis of the metagenomic approaches distinguishing between DIAMOND alignment against the respective protein database and the meganization step for NCBInr-, AnnoTree- or mgPGPT-db mappings.

Regarding the detection of entirely 5,634 PGPTs (level 6), 5,230 traits were detected by all metagenomic PGPT approaches (Figure 4 D). 159 PGPTs were only detected by mgPGPT-, 12 only by AnnoTree-, and 11 only by NCBInr-db based analyses. 15 additional PGPTs were annotated when applying either NCBInr- or AnnoTree-db. However, the analysis applying the mgPGPT-db is able to recover most of the traits, namely 5,596.

The runtime of the mgPGPT-db diamond and meganization runs of 35 samples are the lowest followed by the AnnoTree-db based analysis (Figure 4 E). By far the NCBInr-db alignment and consecutive meganization steps are most time consuming with runtime 2.6 times higher than applying AnnoTree-db and 7.4 times higher than mgPGPT-db (Table S3).

### Results of the MWAS study comparing PGPT abundancies across environments

The metagenome-wide association of 35 samples has be considered not only as a performence test but also a proof of concept if the PGP-specific patterns can be achieved by mgPGPT based metagnomic analysis. Indeed, on PGPT ontology level 2 for instance PGPTs related to biofertilization, plant immune response stimulation or stress control attribute to plant associated samples (Figure S1 A). In contrast the PGPT classes competitive exclusion and colonizing plant systems show higher enrichment regarding animal and human association. These enrichments can be directly associated to functional orthologous PGPT protein groups on level 6 (Figure S1 B), where a better clustering of the samples into plant+soil and Human+animal can be reproduced. Respective PGPT identifiers and their abundance values can be taken from Table S4.

## DISCUSSION

The annotation of PGPTs in metagenomic samples opens a new perspective for plant metagenome analysis. While in the past the description of PGPTs in genomes and metagenomes was based on individual gene selection and not standardized across studies, the PGPT ontology implementation in MEGAN6 enables entire PGPT functional analysis of short and long reads according the DIAMOND+MEGAN pipeline (28). Already NCBInr-db or AnnoTree-db based annotated metagenomes can now be easily re-meganized, using their respective mapping databases, and analysed with the MEGAN PGPT viewer, what avoids the time- and memory-extensive realignment. Alternatively, metagenomic samples might be aligned with DIAMOND directly against mgPGPT-db, an optimized database comprising metagenomic protein sequences retrieved from AnnoTree, including proteins e.g., curated from MAGs (34, 35). Not surprisingly, the read assignment rate of mgPGPT-db, the MEGAN NCBInr-db and AnnoTree-db based PGPT mapping approximate each other and are highly recommended for use when dealing with soil or plant samples, where the portion of yet unknown microbial content is still enormous (35). If possible, long read sequencing data achieved by e.g., Oxford Nanopore or PacBio should be preferred, as it provides the opportunity to detect whole syntenic PGPT gene clusters along a read and offers the opportunity to assembly complete genomes including chromosomes and plasmids as MAGs (37). That can lead to the identification of entire functional syntenic gene clusters, like biosynthetic gene clusters (BGCs) in metagenomes (38). Means, when using short reads you might miss genomic structural PGPT information of single organisms, albeit assembling the genomes, what even does not appear very accurate on short reads except a very high sequencing depth by low host contamination (39).

The performance of metagenomic functional PGPT-based clustering is highly comparable between all approaches. The outliers detected must be addressed in future by e.g., taking at least 4 samples per different within-group sites and entirely more metagenomic samples. Here the small-scale MWAS study was just used as proof of concept and not to get most abundant plant-associated PGPTs. However, we could identify enriched plant-associated genes, related to e.g., biofertilization, plant immune respone stimulating agents, stress control or other putative traits that were also reported by Levy and coworker (40). An increased PGPT value for human and animal associated bacterial communities regarding the classes competitive exclusion and colonizing the plant system suggest the importance of such genes not only in the plant microbiome but much higher in other host systems (e.g., animal and human) for best persistence. Interestingly, the PGPT ontology seems to be able to distinguish between rhizospheric from phyllospheric communities – but to confirm that, more data must be studied and the plant species and geographic location should be taken into account to a greater extent. Further, our analysis points to overlapping PGPT traits pattern between most rhizospheric and soil-associated samples, what might be explained by the steady exchange between respective microbial communities across their shared environment (41). The close PGPT-based clustering of the sole human gut sample with the plant samples have to be proved in detail with more samples in future to maybe conclude to specific human nutrition behaviours (42). Comparing the PGPT annotation rate among all PGPT annotation strategies, we have to mention that for all unique annotated PGPTs appear. But among the metagenomic approaches their count is quite small. Regarding the point of an enriched mgPGPT-db including proteins from MAGs, that are not yet part of the PGPT-db, we are about to also provide the mgPGPT-db for PGPT-Pred on PLaBAse for genome annotation in the near future.

When considering all arguments, applying the mgPGPT-db approach might be the preferable option to go for if you like to explore the PGP potential of plant or soil metagenomes. Aside the facts that more PGPT protein sequences can be assigned resulting in a higher read assignment rate, the runtime is lower compared to NCBInrdb and AnnoTree-db protein alignments, due to the alignment against a smaller non-redundant database that reduces diamond alignment and meganization time. However, if you are interested in both, entire functional annotations, that are provided by e.g., the EGGNOG, INTERPRO2GO, KEGG or SEED ontologies, and PGPTs, you might go for either the AnnoTree or the NCBInr approach. That is because the mgPGTP-db comprises only a subset of proteins compared to these databases. As the taxonomic and functional affiliations show best comparability between mgPGPT-db and AnnoTree-db analyses and the run time is still adequate, one could prefer the AnnoTree-db based analysis for a holistic approach.

Altogether, we consider the DIAMOND-MEGAN-mgPGPT approach a significant improvement for future investigations of the bacterial PGPT potential, especially of plant and soil metagenomes, but also of other communities. It will contribute the understanding of the PGPT variation dependent on e.g., the surrounding biotic and abiotic conditions or host genotypes. Additionally, respective taxonomic remapping of PGPTs by the MEGAN tool *taxonomy2function* might lead to identification of high PGPT-containing strains, such as plant growth-promoting bacteria (PGPBs), and might facilitate the selection of promising, site-specific applicable bioinoculant strains. Exemplarily, comparing the PGPT content of natural occurring microbiomes of stress-susceptible versus stress-resistant plants would allow the exact definition of PGPTs involved and their association among the bacterial community, what assists even the definition of synthetic synergisitc communities (SynComs) (43). Additionally, the standardized metagenomic PGPT annotation approach also guaranties better comparability between different studies, microbial communities, environments, and single strains around the world. Moreover, the integration of the PGPT ontology into other metagenome analysis tools, like e.g., in MAIRA for real-time and/or in field microbial tracking, MMseqs2, and BugSplit or into (meta-) transcriptomic analysis via gene set enrichments are conceivable (44, 45, 46).

## MATERIALS AND METHODS

All scripts mentioned in the following subsections are available via the git repository mgPGPT.

### PGPT ontology implementation in MEGAN6

The PGPT ontology, published by Patz and coworkers (36) was implemented into MEGAN 6 by converting it into the MEGAN-specific tree (.tree) and map file (.map). For the hierarchical tree file generation each internal node name in the PGPT ontology was replaced by a unique, numeric node identifier. The .map file lists the respective node names to each node id, so that the entire ontology can be rebuild in the MEGAN graphical user interface (GUI) as PGPT-Viewer. A respective button is added to the menu bar. At leaf level, PGPT identifier might appear multiple times according to sharing functions across the ontology, means the tree is multi-labeled. A PGPT identifier represents a functional orthologous group. Consequently a PGPT identifier is associated to multiple proteins in the respective protein database.

### mgPGPT protein sequence database curation

The mgPGPT protein database, mgPGPT-db, was achieved by extending the current PGPT-db (36), by protein sequences stored in the Megan-AnnoTree-db (35), that share PGPT specific K numbers, but are absent in the PGPT-db. Therefore, respective sequences of both databases were hashed and sequences with non-overlapping hash values considered extension only.

In a next step, a non-redundant sequence list was curated with uclust v1.2.22q:

~~~
uclust --sort <at-PGPTid-prot.faa> --output <at-PGPTid-prot_sor.faa>
uclust --input <at-PGPTid-prot_sor.faa> --uc at-PGPTid-prot_res.uc> --id 1.00 --amino
uclust --uc2fasta <at-PGPTid-prot_res.uc> --input <at-PGPTid-prot_sor.faa> --output <at-PGPTid-prot_non-red.faa> --types
~~~

Non-redundant sequences were integrated into the PGPT-db forming the new mgPGPT-db. The header of added sequences was replaced by the PGPT identifier and a consecutive number, thus both together building up a unique sequence identifier.

Finally, a mgPGPT protein database for (i) blast and for (ii) diamond for gene, protein or metagenomic short and long read alignments in blastx or blastp mode, respecitively, were created:

~~~
makeblastdb -in <PGPTid-proteins_non-red.faa> -title mgPGPT-db -parse_seqids -dbtype prot
diamond makedb --in <PGPTid-proteins_non-red.faa> -d <mgPGPT_APR2022-db.dmnd>
~~~

### Setting up the MEGAN6 mgPGPT mapping database

The mgPGPT mapping database file for MEGAN, was generated with SQLite3 (version 3.32.3):

~~~
sqlite3 megan-map-mgpgpt-APR2022.db
sqlite$ CREATE TABLE info (id TEXT PRIMARY KEY, info_string TEXT, size NUMERIC);
sqlite$ CREATE TABLE mappings (Accession PRIMARY KEY, Taxonomy INT, GTDB INT, PGPT INT) WITHOUT ROWID;
~~~

The mgPGPT mapping database contains two tables. The info table lists the content and versions for entries of the mapping database. The mappings table provides MEGAN node ids for taxonomic and PGPT functional annotations to every PGPT protein accession (unique key) present in the mgPGPT blast and diamond database files. The taxonomic affiliations of each protein sequence were fetched from the NCBInr and Annotree mapping database entries for NCBI and GTDB taxonomies (megan-map-Feb2022-ue.db), based on the hash sequence comparison results. In case of remaining non-identical sequences taxonomic affiliation was achieved by aligning against the NCBI nr protein database with DIAMOND and taxonomic mapping by applying the LCA algorithm, considering top 10 hits:

~~~
diamond blastp -q <remain_PGPTproteins.fasta> --db <nr-proteins.dmnd> -o <remain_PGPTproteins.daa> -f 100 –fast
daa-meganizer -i <remain_PGPTproteins.daa> -mdb <NCBInr_mappingDB> -alg naive -top 10 -me 1e-5 -mpi 97
~~~

Afterwards, the MEGAN tool daa2info provided NCBI taxonomy node ids for each PGPT protein:

~~~
daa2info -i <remain_PGPTproteins.daa> -o <remain_PGPTproteins2tax.txt> -r2c Taxonomy
~~~

For a few cases of 0,0025% of the PGPT sequences no taxonomic rank could be assigned. Their respective taxonomic database entries remained unassigned.

During meganization the option -mdb allows loading the generated mgPGPT mapping database file.

### PGPT extension for MEGAN6 NCBInr and AnnoTree mapping databases

As an alternative, the conservative DIAMOND+MEGAN approach via alignment against NCBInr or AnnoTree mapping database, can be applied. Therefore, by hashing all sequences as described, we computed the overlap between sequences listed in the mgPGPT protein database and all above mentioned databases, respectively. Based on the resulting shared sequence hashes we added the associated PGPT-ontology node id to the NCBInr or AnnoTree mapping database into a new column, called PGPT:

~~~
sqlite3 megan-mapping.db
sqlite$ ALTER TABLE mappings ADD PGPT INT
sqlite$ INSERT INTO info (id, info_string, size) VALUES (“PGPT”,”created: Thu Feb 17 11:17:00 CET 2022 Cite: Patz et al 2022)”,
sqlite$ update_mappingDB_sql3.py <megan-mapping> <NCBInrProtAcc2PGPTnodeID.txt>
~~~

mgPGPT mappings are available for NCBInr and AnnoTree mapping database releases.

All files needed for the diamond+MEGAN6 mgPGPT-targeted metagenome analysis can be downloaded from the PLaBAse web resource.

### Metagenomic dataset curation

For metagenomic application of the PGPT ontology, 35 random public available Illumina short read data sets, comprising 14 plant-associated (PA), 14 non-plant-associated (NPA) and 7 soil metagenomes, were downloaded from the MG-RAST server (47). PA samples can be further distinguished between 7 rhizospheric and 7 phyllospheric samples. NPA samples can be associated to two environmental groups: human- (HA) and animal-associated (AA). Human samples were mainly acquired from the Human Microbiome Project (HMP) (48). Respective classifications were given by the MG-Rast metadata collection per sample that follows the Genomic Standard Consortium (GSC) metadata standards (49). All data files analyzed with our pipeline had passed already the MG-RAST quality checks and filters. We applied seqkit to obtain a read statitic per sample, what results in read of average 100-200 bp. All samples did reach 80% of bacterial assigned reads and a bacterial read count of at least 500,000 based on taxonomic analysis according to the DIAMOND-MEGAN6 pipeline (30, 28, 50). Their accession numbers, metainformation, respective sample class, and read statistics are listed in Table S5. For metagenome analysis, of all samples only reads assigned to the domain Bacteria (-n) the taxonomic (-c) ranks below (-p) were extracted:

~~~
‘
read-extractor -i <sample.daa> -c Taxonomy -n Bacteria -b true
~~~

### Metagenomic Analysis

According to the DIAMOND-MEGAN6 pipeline (28) reads of all samples were aligned against either the NCBInr, AnnoTree, PGPT and mgPGPT protein database with DIAMOND v2.0.9.147 24. DIAMOND ran on each metagenomic sample as follows:

~~~
diamond blastx --db <database.dmnd> --query <sample_bact.fasta.gz> --out <sample> --outfmt 100 -b5 -c 1 -e 1e-5
~~~

The resulting alignments were saved as DIAMOND alignment archive files (e.g. sample.daa) and mapped with either the NCBInr (previous/recent), AnnoTree or PGPT (mg/PGPT-db) mappings database using MEGAN daa-meganizer:

~~~
daa-meganizer -i <sample.daa> -mrc 90 -mpi 60 -mdb <mapping-database.db>
~~~

Single samples were then further analyzed with the MEGAN GUI as described for the DIAMOND-MEGAN6 pipeline.

### Metagenome-wide association study

A metagenome-wide association study (MWAS) of PGPT was performed for all samples in association to their environmental classes per each approach. Firstly, the read counts of all samples were normalized (-n true) to the lowest read count, found in any sample with the MEGAN comparison tool:

~~~
compute-comparison -i <allSamplesDAA-Folder> -o <comparison.megan> -n <true/false> <-k1 true>
~~~

Non-zero read count values caused by normalization of very low red counts were mapped to at least one counted read (-k1 true) to achieve best comparability of e.g., detected PGPT ids by all methods.

The normalized samples were then grouped and colored according their environmental groups within the Sample viewer, after loading the respective metadata file. The beta diversity among all samples was then calculated and demonstrated as PCoA plots based on Bray Curtis ecological index within the MEGAN GUI to cluster all samples according their PGPT abundance values and taxonomy. Likewise, repsective outline graphs were drawn.

### Performance Analysis of MEGAN PGPT Annotation Strategies on metagenomic samples

For comparison of the four approaches, the PGPT read alignment and assignment rate in total, as well as on PGPT ontology level 1 were calculated for all single metagenomic samples and presented for one sample or as a sum of all samples. We run the comparison mentioned above again but with non-normalized raw read counts (-n false). As another quality measurement the amount of annotated PGPT ids and PGPT protein overlapps of all approaches were investigated and visualized with the R package venn. Additionally, the runtime for the entire MWAS study was traced for DIAMOND alignment against the NCBInr, AnnoTree, PGPT-db, and mgPGPT-db protein databases and for the consecutive meganization step of DAA files.

### Hardware specification

The entire analysis was performed on BwForCluster BinAC, where each node comes up with a 2x Intel Xeon E5-2630v4 (Broadwell) 2.4 GHz processors with 28 cores and maximal 128 GB working memory.

### Data availability

Analyzed datasets are open source and available via the MG-RAST Server and listed in Table S5. All exploited data and generated protein and mapping databases locations are publicly accessible via the MEGAN releases from May2022 on and our websites, specified in its respective method section above.

## SUPPLEMENTAL MATERIAL

**Note**; All supplemental data, that is listed below, will be made available on publication.

**Figure S1**. Metagenome-wide association study comprising five environmental classes performed with the mgPGPT-db approach.

All supplemental tables are available in the file mgPGPT_SupplementalTables.xlsx.

**TABLE S1**. Shared PGPT functional and taxonomic assignment rates for PGPT-assigned reads only, when analyzing the single metagenomic dataset mgm4626743.3 with all three mgPGPT approaches.

**TABLE S2**. Runtime analysis applying three metagenomic PGPT annotation approaches compared to the recent PGPT-db for sample mgm4626743.3.

**TABLE S3**. MWAS runtime analysis applying three metagenomic PGPT annotation approaches compared to the recent PGPT-db.

**TABLE S4**. mgPGPT-db based MWAS study comparing PGPT abundancies for the classes plant- (PA), human- (HA), animal- (AA), soil-associated (SA) and others on PGPT ontology level 6.

**TABLE S5**. Metagenomic data set description.

## ACKNOWLEDGMENTS

We acknowledge the state Baden-Wuertemberg (Germany) for providing the bwFor-Cluster BinAC Server capacities for high-throughput computing and the Julius Kühn Institute - Federal Research Centre for Cultivated Plants in Braunschweig (Germany) for its contribution to the common hardware deployment. Further, we would also like to acknowledge the support by the BMBF-funded de.NBI Cloud within the German Network for Bioinformatics Infrastructure (de.NBI) (031A532B, 031A533A, 031A533B, 031A534A, 031A535A, 031A537A, 031A537B, 031A537C, 031A537D, 031A538A) for this work.

## ABBREVIATIONS

BGCs: biosynthetic gene clusters
db: database
MAG(s): metagenome assembled genome(s)
MAIRA: mobile analysis of long reads
MEGAN: metagenome analyzer
MWAS: metagenomic association study
NCBI: National Center for Biotechnology Information
nr: non-redundant protein database
mg: MEGAN / metagenomic
PGP: plant growth-promoting / plant growth-promotion
PGPB: plant growth-promoting bacteria
PGPT(s): plant growth-promoting gene(s) / protein(s) / trait(s)
PLaBAse: plant-associated bacteria web resource
Pred: prediction tool

